# Prophages and plasmids display opposite trends in the types of accessory genes they carry

**DOI:** 10.1101/2022.07.21.500938

**Authors:** Nobuto Takeuchi, Sophia Hamada-Zhu, Haruo Suzuki

**Affiliations:** School of Biological Sciences, the University of Auckland, Private Bag 92019, Auckland 1142, New Zealand, Universal Biology Institute, the University of Tokyo, 7-3-1 Hongo, Bunkyo-ku, Tokyo 113-0033, Japan; School of Biological Sciences, the University of Auckland, Private Bag 92019, Auckland 1142, New Zealand; Institute for Advanced Biosciences, Keio University, Tsuruoka, Yamagata, Japan, Faculty of Environment and Information Studies, Keio University, Fujisawa, Japan

**Keywords:** virus, bacteriophage, selfish genetic elements, evolution of pathogenicity, evolution of antimicrobial resistance, lateral gene transfer

## Abstract

Mobile genetic elements (MGEs), such as phages and plasmids, often possess accessory genes encoding bacterial functions, facilitating bacterial evolution. Are there evolutionary rules governing the arsenal of accessory genes MGEs carry? If such rules exist, they might be reflected in the types of accessory genes different MGEs carry. To test this hypothesis, we compare prophages and plasmids with respect to the frequencies at which they carry antibiotic resistance genes (ARGs) and virulence factor genes (VFGs) in the genomes of 21 pathogenic bacterial species using public databases. Our results indicate that prophages tend to carry VFGs more frequently than ARGs in three species, whereas plasmids tend to carry ARGs more frequently than VFGs in nine species. In *Escherichia coli*, where these trends are detected, prophage-borne VFGs encode a much narrower range of functions than do plasmid-borne VFGs, typically involved in damaging host cells or modulating host immunity. In the species where the above trends were not detected, ARGs and VFGs are barely found in prophages and plasmids. These results indicate that MGEs differentiate in the types of accessory genes they carry depending on their infection strategies, suggesting an evolutionary rule governing horizontal gene transfer mediated by MGEs.

## 3. Introduction

Mobile genetic elements (MGEs), such as phages and plasmids, often possess accessory genes encoding bacterial phenotypes that are not necessarily integral to the replication or transmission of MGEs. For example, plasmids frequently possess various antibiotic resistance genes (ARGs) [1–3]. Also, phages and plasmids possess virulence factor genes (VFGs) required for bacterial pathogenicity [4–8]. Mediating the horizontal transfer of accessory genes, MGEs play important roles in the evolution of bacterial genomes and phenotypes [9, 10].

MGEs are parasites of bacteria. Thus, horizontal gene transfer (HGT) mediated by MGEs can be regarded as the genetic manipulation of hosts by parasites [11]. Given that MGEs are self-interested evolving entities, MGEs are expected to possess accessory genes that advantage themselves [12, 13]. For example, plasmids are considered to gain selective advantages from ARGs by improving the survival of their bacterial hosts in heterogeneous environments [13–16]. It has also been hypothesised that phages gain selective advantages from VFGs by modifying environments in which their bacterial hosts live [17].

What evolutionary rules govern the arsenal of accessory genes carried by MGEs [12, 13]? Such rules, if they exist, might reflect the different infection strategies of MGEs. For example, phages typically lyse host cells to transmit to other cells, whereas plasmids do not. Consequently, phages might not gain much of an advantage by carrying genes that improve the survival of bacteria, such as ARGs, because if their hosts are in danger, they can enter a lytic replication cycle to abandon their hosts and seek new ones [18–22]. By contrast, plasmids cannot typically use such evacuative strategies, hence likely to benefit from genes improving host survival. Thus, to understand the rules governing HGT mediated by MGEs, it is beneficial to investigate whether different MGEs carry different types of accessory genes.

To address the above question, we consider an ongoing debate about phage-borne ARGs. While it is well established that plasmids frequently carry ARGs [1–3], how frequently phages carry ARGs is controversial [23]. Phages mediate HGT through multiple mechanisms, among which specialised transduction is the most similar to HGT mediated by plasmid [24] (see Discussion for the other mechanisms). In specialised transduction, phages transfer genes carried in their genomes. Therefore, specialised transduction is strictly coupled with the infectious transmission of phage genomes, the coupling that is also entailed in plasmid conjugation [24]. Laboratory experiments have demonstrated that phages are capable of transferring ARGs to bacteria through specialised transduction [25]. However, the specialised transduction of ARGs in nature has been scarcely documented [3, 26]. While metagenomic studies have detected ARGs in viral fractions of environmental DNA samples [27–30], other studies provide evidence suggesting that the detection of ARGs was due to the contamination of bacterial DNA in the viral fractions [31, 32]. Genomics studies have predicted a number of prophages—i.e., phage genomes inserted into bacterial chromosomes as a consequence of specialised transduction—carrying ARGs in the genomes of *Acinetobacter baumannii* [33], *Klebsiella pneumoniae*, and *Pseudomonas aeruginosa* [34] (see also [35]). Also, a previous study has isolated 29 phages from wastewater, of which 15 carry ARGs, suggesting that phages frequently possess ARGs [36]. However, these results appear at odds with a recent comprehensive analysis of phage genomes in public databases, which shows that ARGs are carried by only 0.3% of phages [37]. Taken together, the existing studies present mixed messages about the frequency at which phages carry ARGs.

To investigate how frequently phages carry ARGs, here we compare the distributions of ARGs and VFGs between the prophages and plasmids of pathogenic bacteria by comprehensively analysing public databases. We consider prophages instead of phages to compare different MGEs belonging to the same bacterial genomes. Our approach is designed to mitigate two issues we consider to be involved in the computational analyses of ARGs encoded in prophages, which are not taken into account in previous studies [33–35]. First, the misidentification of prophages can cause systematic biases in the number of prophage-borne ARGs. For example, non-prophage regions can be misidentified as prophages, causing an overestimation in the number of prophage-borne ARGs. Contrariwise, a true prophage can be missed, which leads to an underestimation of the number of prophage-borne ARGs. To avoid these biases due to prophage prediction, we compare the number of prophage-borne ARGs to that of prophage-borne VFGs, where both numbers are expected to be biased by common factors so that the biases can be cancelled out. The second issue involved in the analysis of prophage-borne ARGs is a sampling bias in bacterial genomes, which can cause overestimation in the numbers of ARGs and VFGs owing to the double-counting of orthologous genes. The degree to which this bias occurs can differ between ARGs and VFGs. To correct this bias, we cluster all genes into putative orthologous groups based on sequence similarity and synteny conservation and count the numbers of putative orthologous groups of ARGs and VFGs (OGARGs and OGVFGs, respectively). Furthermore, to investigate a potential differentiation between prophages and plasmids, we analyse the distributions of ARGs and VFGs in plasmids. Finally, we examine whether prophages and plasmids also differ in the functional categories of VFGs they carry. Our results suggest that prophages are biased towards carrying VFGs, whereas plasmids are biased towards carrying ARGs. However, in many species, both ARGs and VFGs are hardly detected in prophages and plasmids. Moreover, we found that prophage-borne VFGs are less functionally diverse than plasmid-borne VFGs in *Escherichia coli*. Taken together, these results indicate that prophages and plasmids differ in the types of accessory genes they carry.

## 4. Methods

Our method is sketched in Figure 1.

**Figure 1.**
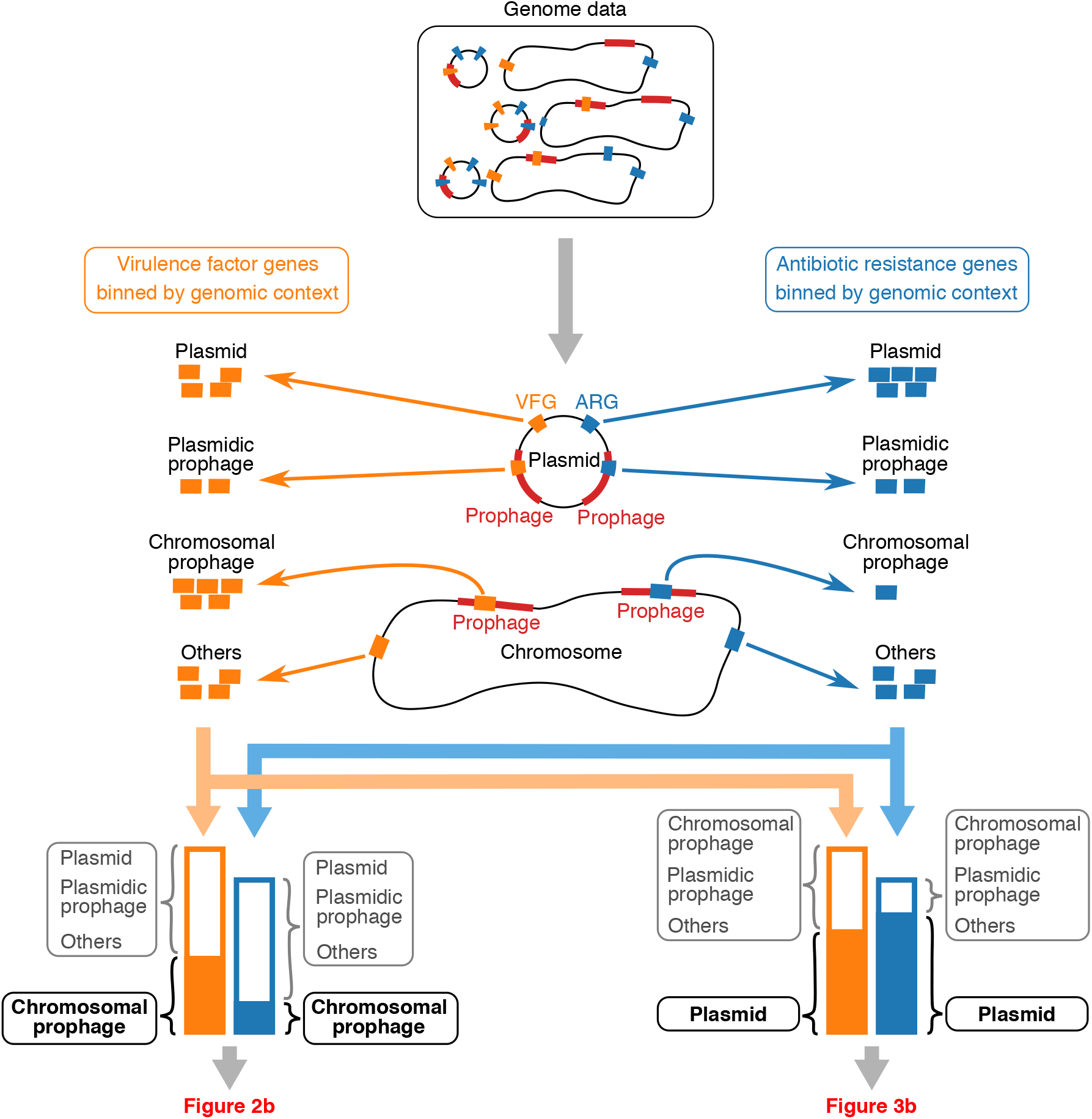
Schematic drawing of the method used in this study. For simplicity, clustering of orthologous genes is omitted from depiction (see Methods).

### 4.1 Data acquisition

Three VFG databases, namely, VFDB (3685 genes in set A), Victors (5085 genes), and PATRIC_VF (1293 genes), were downloaded from the respective websites in December 2020 [38–40]. The VFG entries in Victors were refined by removing those carried by non-bacterial pathogens or lacking NCBI protein GIs (4575 out of 5085 remained). Some VFG entries in Victors were missing protein sequences, which were downloaded from Genbank based on their protein GIs [41]. All VFG entries in the three databases were pooled and clustered to remove redundancy with CD-HIT with the protein sequence identity threshold of 1.0 [42], resulting in a combined VFG database of 7218 entries.

The genome assemblies with the ‘Complete’ status were downloaded from RefSeq in September 2021 with the following criteria [43]: a species had at least 60 complete genomes in RefSeq and at least 70 VFGs in the combined VFG database. These criteria resulted in 21 species of bacterial pathogens spanning three phyla, Actinobacteria, Firmicute, and Proteobacteria, as follows (numbers in brackets indicate the number of genomes examined in this study): *Acinetobacter baumannii* (275), *Bacillus anthracis* (99), *Bordetella pertussis* (562), *Brucella melitensis* (64), *Burkholderia pseudomallei* (126), *Campylobacter jejuni* (220), *Enterococcus faecalis* (64), *E. coli* (1444; *E. coli* was considered to include *Shigella boydii* (17), *Shigella dysenteriae* (21), *Shigella flexneri* (35), and *Shigella sonnei* (1)), *Haemophilus influenzae* (92), *Helicobacter pylori* (225), *Klebsiella pneumoniae* (873), *Legionella pneumophila* (102), *Listeria monocytogenes* (263), *Mycobacterium tuberculosis* (285), *Neisseria meningitidis* (122), *Pseudomonas aeruginosa* (320), *Salmonella enterica* (996), *Staphylococcus aureus* (618), *Streptococcus agalactiae* (91), *Streptococcus pyogenes* (235), and *Vibrio cholerae* (99). The complete list of 7175 genomes analysed in this study is in Table S1.

### 4.2 Prophage prediction

To predict prophages in the bacterial genomes, VIBRANT (version 1.2.1) was used for the following reasons. First, VIBRANT has comparatively high performance, as reported by a recent benchmark [44]. Second, it is a standalone tool, so it can be run on local computers. Third, its algorithm is based on the similarity search of known phage proteins rather than predicting prophages based on nucleotide signatures. VIBRANT was run against the genomic sequences with default parameters.

### 4.3 Plasmid prediction

Sequences (contigs) were considered as plasmids or chromosomes if they were annotated as such in the RefSeq assembly report files (in total, there were 9279 plasmid and 6607 chromosome contigs). Contigs annotated as ‘Segment’, ‘Genome Segment’, or ‘Extrachromosomal Element’ were ignored (3, 1, or 5 contigs, respectively).

### 4.4 ARG prediction

To predict ARGs in the bacterial genomes, AMRFinderPlus (version 3.10.5) was run against translated coding sequences with the core subset of the database (version 2021-09-11.1) and, if possible, the organism option enabled [45]. The genes predicted by AMRFinderPlus as ARGs (i.e., ‘element subtype AMR’) and not annotated as pseudo-genes in RefSeq were considered as ARGs (those predicted as ‘element subtype POINT’, which contain point mutations associated with AR, were excluded).

### 4.5 VFG prediction

To predict VFGs in the bacterial genomes, every entry in the combined VFG database (VFG database entry, for short) was queried against the translated coding sequences of every bacterial genome with BLASTP with an E-value threshold of 1e-9 [46]. A gene in a bacterial genome (bacterial gene, for short) could match multiple VFG database entries, in which case the VFG database entry with the highest bit-score was selected as the best match. A bacterial gene was considered as encoding VF if it met the following additional criteria: (i) it was not annotated as a pseudo-gene in RefSeq [43]; (ii) the BLASTP alignment between the bacterial gene and its best-match VFG database entry, if any, had at least 80% sequence identity and covers at least 80% of both the bacterial gene and the best-match VFG database entry; (iii) the species of the genome in which the bacterial gene resides was identical to the species in which the best-match VFG database entry resides [38–40].

### 4.6 Orthology prediction

To avoid double-counting orthologous genes in closely related strains, all genes within a species were clustered into putative orthologous groups based on protein sequence similarity and synteny conservation, as follows. First, preliminary orthologous pairs of genes were identified between every pair of genomes within each species through all-against-all sequence similarity searches using ProteinOrtho version 6.0.25 (with DIAMOND ver. 2.0.6 [47]; E-value cut-off of 1e-5; minimum coverage of best alignments of 75%; minimum percent identity of best alignments of 25%; minimum reciprocal similarity of 0.95) [48]. ProteinOrtho defines a preliminary orthologous pair of genes as a reciprocal nearly-best hit (RNBH), as follows. A nearly-best hit (NBH) of a gene queried against a target genome is defined as a hit whose bit-score is not smaller by a factor *f* than that of the best hit. The value of *f* was 0.95, which is the default value of ProteinOrtho. If two genes are mutually NBH of each other, they form RNBH [48].

RNBHs obtained with ProteinOrtho were pruned based on synteny conservation with an in-house script, as follows. Let *x* and *y* be a pair of genes forming RNBH, and let *X* and *Y* be the genomic neighbours of *x* and *y*, respectively, where *X* is defined as a set of 21 genes consisting of ten genes upstream of *x*, ten genes downstream of *x*, and *x* itself, and *Y* is likewise defined in terms of *y* (all contigs were assumed to be circular, and the orientation of genes were ignored). Let *N_x_* and *N_y_* be the number of genes in *X* and *Y* that form RNBHs with at least one gene in *Y* and *X*, respectively (note that a single gene in one genome can form RNBHs with multiple genes in another genome owing to tandem duplication). If both *N_x_* and *N_y_* are greater than ten (i.e., a majority of the genes in *X* form RNBHs with the genes in *Y*, and *vice versa*), the RNBH formed by *x* and *y* was kept; otherwise, it was discarded [49].

Finally, the pruned RNBHs were clustered into putative orthologous groups with the spectral clustering algorithm implemented in ProteinOrtho version 6.0.25 (minimum algebraic connectivity of 0.1; exact step 3; minimum number of species of 0; purity of 1e-7) [48].

### 4.7 Classification of orthologous gene groups

A gene (VFG or ARG) was considered to be encoded in a prophage if the entire gene is included within a genomic region predicted as a prophage. Similarly, a gene was considered to be encoded in a plasmid if the gene resides in a plasmid contigue. Moreover, genes in prophages residing in plasmids (plasmidic prophages, for short) are distinguished from those in prophages residing in chromosomes (chromosomal prophages) for two reasons. First, it was ambiguous whether genes in plasmidic prophages should be regarded as encoded by plasmids, prophages, or both. Second, plasmidic prophages potentially represent a distinct class of MGEs called phage-plasmids [50].

An orthologous group of genes was considered to be encoded in chromosomal prophages, plasmidic prophages, or plasmids if the majority of the genes belonging to the group were encoded in the respective genomic contexts (the cases of ties were ignored).

An orthologous group of genes was considered an ARG or VFG if the majority of the genes belonging to the group were predicted as ARGs or VFGs, respectively. The majority rule was used because a subset of genes in OGARG or OGVFG could be predicted as non-ARGs or non-VFGs, respectively, owing to sequence divergence. However, for most orthologous groups of ARGs and VFGs, all genes in a group were predicted as either ARGs or VFGs. Moreover, no orthologous group contained both ARGs and VFGs.

### 4.8 Functional classification of prophage- and plasmid-borne VFGs

To classify the functions of VFGs, gene symbols (i.e., abbreviated gene names) were assigned to prophage-borne and plasmid-borne OGVFGs in *E. coli*, as follows. All OGVFGs in *E. coli* genomes were collectively associated with 862 best-match VFG database entries (see Method under ‘VFG prediction’). Each of these entries was looked up in the respective VFG database (VFDB, Victors, or PATRIC_VF) to find a gene symbol assigned to it [38–40]. When a gene symbol was missing, a gene symbol was manually determined by querying the protein sequence of a VFG database entry against VFDB, the NCBI Conserved Domain Database (CDD), and the NCBI NR database [38, 51, 52] (Table S2). Consequently, 856 out of the 862 VFG database entries were assigned gene symbols (Table S2). The remaining six entries, which are annotated as hypothetical proteins in the VFG databases, were discarded from the subsequent analyses together with the OGVFGs associated with them. Next, prophage-borne and plasmid-borne OGVFGs were assigned gene symbols based on what VFG database entries they were associated with. If an OGVFG was associated with VFG database entries having an identical gene symbol, that gene symbol was assigned. If an OGVFG was associated with multiple VFG database entries with conflicting gene symbols (only 17 out of 3922 OGVFGs were of this type), the conflict was manually resolved by querying these entries against VFDB, the NCBI Conserved Domain Database, and the NCBI NR database (Table S2). Consequently, prophage-borne and plasmid-borne OGVFGs were collectively assigned 232 gene symbols (Table S2). Subsequently, these gene symbols were also assigned to OGVFGs that are neither prophage-borne nor plasmid-borne.

The 232 gene symbols were manually categorised by their functions based on information obtained from VFDB, NCBI CDD, and Uniprot [38, 51, 53] (Table S3). While most of the functional categories were adopted from VFDB [38], a new category, ‘phage-related’, was added, and the two categories, ‘exoenzymes’ and ‘enzymes’, were merged into ‘enzymes.’ Six gene symbols were annotated to have multiple functions and, therefore, placed under multiple categories.

## 5. Results

### 5.1 Prophage prediction

To examine the distribution of ARGs in prophages, we computationally predicted prophages using VIBRANT [54] and ARGs using AMRFinderPlus [45] in the genomes of 21 pathogenic bacterial species downloaded from the RefSeq database [43] (Methods; Tables S1 and S4).

To avoid double-counting orthologous ARGs in closely related strains, we clustered all genes within a species into putative orthologous groups based on sequence similarity and synteny conservation (Methods). We then counted the number of orthologous groups of ARGs (OGARGs) encoded in the predicted prophages, distinguishing between prophages residing in bacterial chromosomes and prophages residing in plasmids (chromosomal prophage and plasmidic prophage, respectively, for short). This distinction was made because it was ambiguous whether ARGs in plasmidic prophages should be regarded as encoded in prophages, plasmids, or both (plasmidic prophages potentially represent phage-plasmids, which are a separate class of MGEs from typical phages and plasmids [50]). The result shows that a few to several prophage-borne OGARGs were detected in ten of the 21 examined species (Tables 1 and S5).

**Table 1.**
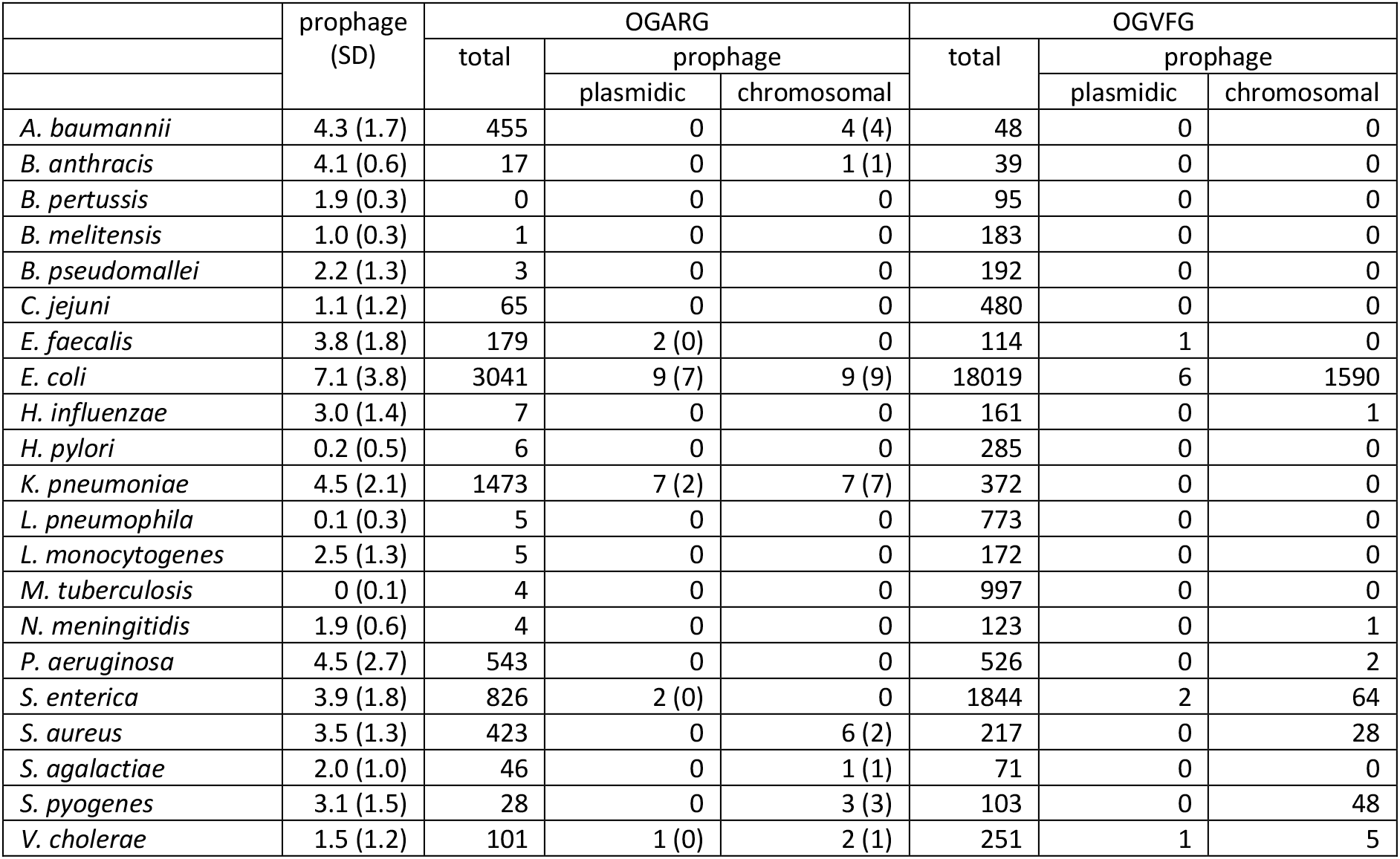
Numbers of predicted prophages per genome (SD, standard deviation), numbers of orthologous groups of antibiotic resistance genes (OGARGs), and those of virulence factor genes (OGVFGs) in genomes (total), predicted plasmidic prophages, and predicted chromosomal prophages. Numbers in brackets are for OGARGs in prophages containing at least one phage-structure gene.

To probe the precision of the above prediction, we manually examined whether predicted prophages carrying ARG contained at least one phage-structure gene (e.g., phage baseplate, capsid, portal, tail, tail fibre, tail sheath, tail assembly, head-tail connector, and tail tape measure) according to the RefSeq annotation. Although this criterion cannot perfectly distinguish true prophages from false ones, it allows us to split the predicted prophages into those enriched with true prophages and those enriched with false prophages, allowing us to probe the precision of the prophage prediction. The result of the examination shows that 24 out of 29 (i.e., 83% of) chromosomal prophages carrying ARGs contained phage-structure genes, whereas seven out of 18 (i.e., 39% of) plasmidic prophages carrying ARGs contained phage-structure genes (Table S6). This result means that 28 out of 33 (i.e., 85% of) OGARGs in chromosomal prophages are in prophages carrying phage-structure genes, whereas nine out of 21 (i.e., 43% of) OGARGs in plasmidic prophages are in prophages carrying phage-structure genes (Table 1; note that the number of prophages carrying ARGs is smaller than that of prophage-borne ARGs because one prophage can carry multiple ARGs). This result suggests that the precision of VIBRANT is acceptable for chromosomal prophages.

During the manual examination, we noticed that a subset of plasmidic prophages carrying ARGs also carried genes encoding integron integrases according to the RefSeq annotation (Tables S6). Moreover, these prophages have a higher frequency of lacking phage-structure genes than those without integron integrase genes (IIGs, for short). Specifically, seven out of ten (i.e., 70% of) prophages carrying both ARGs and IIGs lack phage-structure genes, whereas nine out of 37 (i.e., 24% of) prophages carrying ARGs and lacking IIGs lack phage-structure genes. VIBRANT predicted that the genes annotated by RefSeq as IIGs matched phage integrase genes, which are a typical component of phage genomes. However, the RefSeq annotation implies that proteins encoded by these genes are more similar to integron integrases than phage integrases because the RefSeq annotation considers a broader set of protein families than does VIBRANT [43, 54]. Therefore, the above result suggests that about half of the prophages carrying ARGs and lacking phage-structure genes might have arisen from the misidentification of integrons as prophages (see also [55]).

### 5.2 Prophages carry significantly more VFGs than ARGs in multiple species

To compare the frequencies of ARGs to those of VFGs in chromosomal prophages, we used BLASTP [46] to search bacterial genomes for VFGs collected from VFDB [38], Victors [39] and PATRIC_VF [40] (Methods). We then counted the number of orthologous groups of VFGs (OGVFGs, for short) encoded in the predicted prophages (Tables 1 and S7). The number of OGVFGs cannot directly be compared to that of OGARGs because we have no *a priori* reason to expect that bacteria possess an equal number of VFGs and ARGs. Thus, we instead compared the relative frequencies of OGVFGs and OGARGs against the genomic background (see Figure 1 for the illustration of what we did). Specifically, we performed binomial tests under the null hypothesis that the relative frequencies of OGARGs and OGVFGs in the chromosomal prophages of each species are the same as those of all OGARGs and OGVFGs in the genomes of the respective species. In this test, we included prophages carrying ARGs and lacking phage-structure genes for fairness because prophages carrying VFGs were too numerous to be manually examined (however, we found that none of the prophages carrying VFGs carried IIG, which suggests that the prophages carrying VFGs do not contain false positives arising from integron misidentification). Also, we corrected P values using the Holm–Bonferroni method to control the family-wise error rate of all the statistical tests conducted in this study [56, 57]. The results of the tests indicate that the relative frequencies of OGARGs and OGVFGs in chromosomal prophages are significantly different from those of respective genomic backgrounds in the following three species (Figure 2): *E. coli* (Gammaproteobacteria), *S. enterica* (Gammaproteobacteria), and *S. aureus* (Firmicute). In all these species, chromosomal prophages carry VFGs more frequently than ARGs. The remaining 18 species, where significant biases were not detected, can be grouped into four categories: the one species where many prophage-borne VFGs and no prophage-borne ARGs were detected, but a bias towards VFGs was not statistically significant (*S. pyogenes*),the eight species where only a small number of prophage-borne ARGs or VFGs were detected (*A. baumannii, B. anthracis, H. influenzae, K. pneumoniae, N. meningitidis, P. aeruginosa, S. agalactiae*, and *V. cholerae*), the six species where no prophage-borne ARG and VFG were detected (*B. pertussis, B. melitensis, B. pseudomallei, C. jejuni, E. faecalis, L. monocytogenes*), and the three species where prophages were rarely or hardly detected (*H. pylori, L. pneumophila*, and *M. tuberculosis*) (Table 1 and Figure 2). The paucity or absence of prophage-borne ARGs and VFGs could be due to the limited sensitivity of the prophage prediction tool. However, a large number of prophages were predicted in all species except for *H. pylori, L. pneumophila*, and *M. tuberculosis*. Therefore, if the prophage prediction tool missed prophages carrying ARGs or VFGs, those prophages are likely to be distinct from the currently known phages. Taken together, the above results suggest that prophages tend to carry ARGs less frequently than VFGs if they carry a sufficient number of ARGs or VFGs; however, in many bacterial species, they carry little or no ARGs and VFGs.

**Figure 2.**
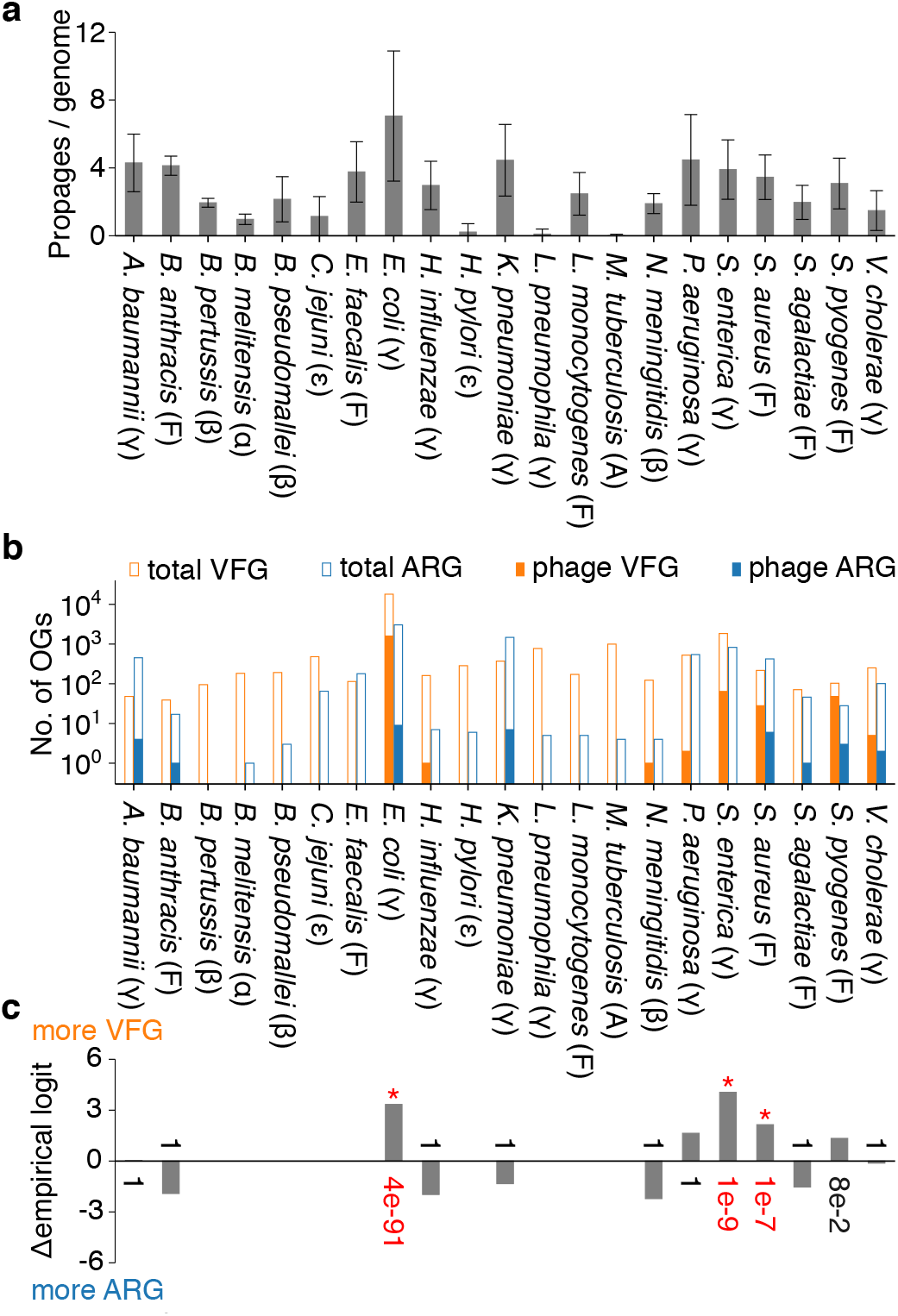
**a** Average number of prophages per genome and standard deviation. **b** Numbers of orthologous groups of virulence factor genes (OGVFGs) and antibiotic resistance genes (OGARGs) in prophages (filled bars) and genomes (open bars). Symbols in brackets indicate taxonomic groups: Actinobacteria (A), Firmicute (F), Alphaproteobacteria (α), Betaproteobacteria (β), Epsilonproteobacteria (ε), and Gammaproteobacteria (γ). **c** Degree of bias towards carrying VFGs more frequently than ARGs. Negative values indicate opposite bias (zero indicates no bias). Difference between empirical logits (denoted as Δempirical logit) is defined as log([*a* + 0.5]/[*b* + 0.5]) – log([*c* + 0.5]/[*d* + 0.5]), where *a* and *b* are numbers of OGVFGs and OGARGs in prophages, respectively (filled bars in b), and *c* and *d* are numbers of all OGVFGs and OGARGs in genomes, respectively (open bars in b). Numbers next to bars are P values of two-sided binomial test under null hypothesis that *a* and *b* are numbers drawn from binomial distribution with probabilities *c*/(*c* + *d*) and *d*/(*c* + *d*), respectively (corrected by Holm–Bonferroni method). Asterisks and red numbers indicate P values less than or equal to 0.05.

### 5.3 Plasmids carry more ARGs than VFGs in many species

To compare prophages to plasmids, we counted the number of OGARGs and OGVFGs encoded in plasmids (Table 2). We then performed similar binomial tests under the null hypothesis that the relative frequencies of OGARGs and OGVFGs in the plasmids of each species are the same as those of all OGARGs and OGVFGs in the genomes of the respective species. The results of the tests indicate that the relative frequencies of OGARGs and OGVFGs in plasmids are significantly different from those of the genomic background in nine out of the 21 examined species (Figure 3). In all nine species, plasmids are biased towards carrying ARGs more frequently than VFGs. These species include six species of Gammaproteobacteria (*A. baumannii, E. coli, K. pneumoniae, P. aeruginosa, S. enterica*, and *V. cholerae*), one species of Epsilonproteobacteria (*C. jejuni*), and two species of Firmicutes (*E. faecalis* and *S. aureus*). The remaining 12 species, where significant biases were not detected, can be grouped into three categories: the four species where plasmids are hardly present *(B. pertussis, B. melitensis, M. tuberculosis*, and *N. meningitidis*), the seven species where plasmids are rare, and plasmid-borne ARGs and VFGs were barely detected (*B. pseudomallei, H. influenzae, H. pylori, L. pneumophila, L. monocytogenes, S. agalactiae*, and *S. pyogenes*), and the one species where plasmids carry more VFGs than ARGs, but a bias was not significant (*B. anthracis*). Taken together, the above results suggest that plasmids tend to carry ARGs more frequently than VFGs if they carry a sufficient number of VFGs or ARGs, with a possible exception of *B. anthracis*; however, in several species, plasmids barely carry VFGs and ARGs.

**Figure 3.**
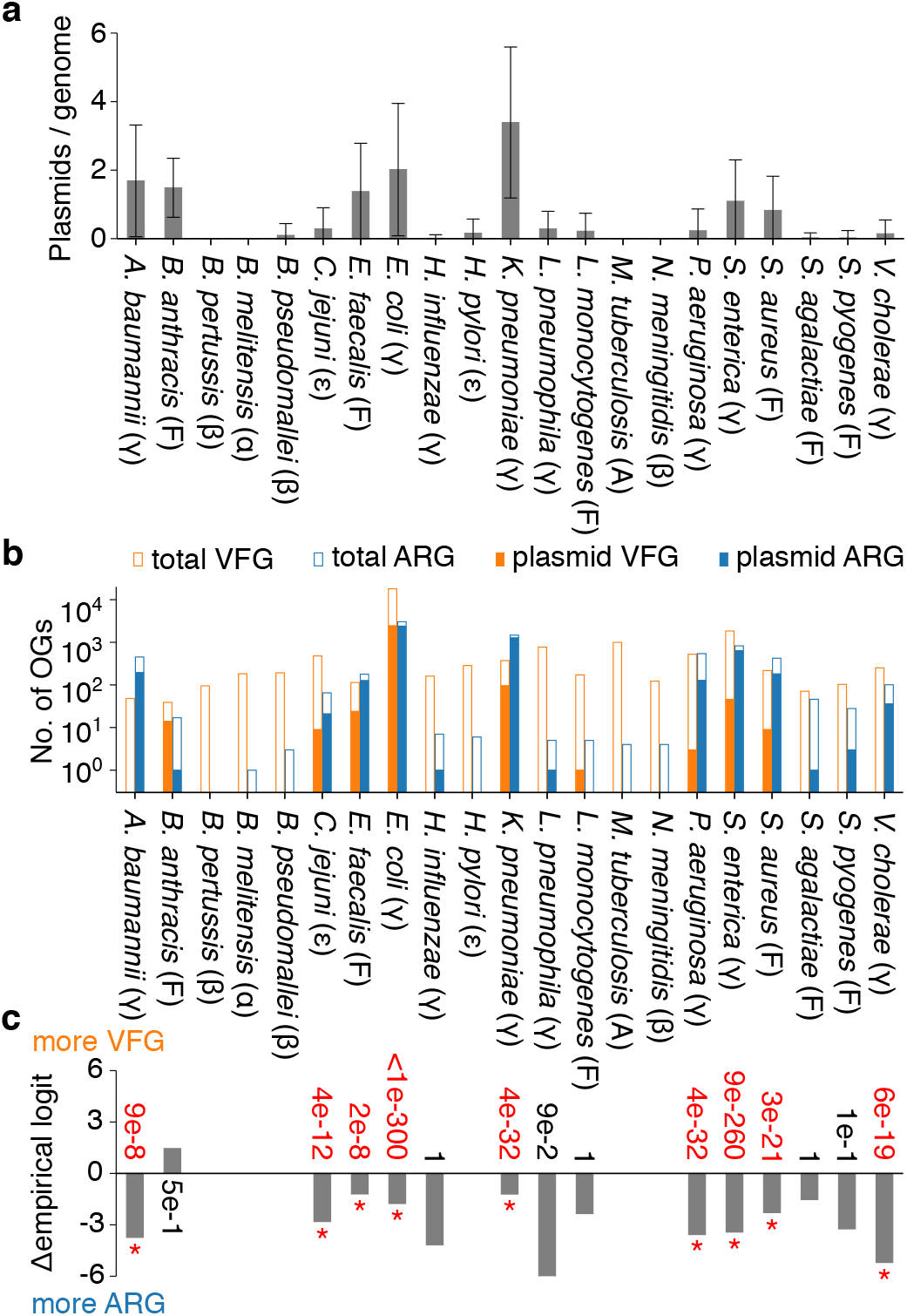
**a** Average number of plasmids per genome and standard deviation. **b** Numbers of orthologous groups of virulence factor genes (OGVFGs) and antibiotic resistance genes (OGARGs) in plasmids (filled bars) and genomes (open bars). Symbols in brackets indicate taxonomic groups: Actinobacteria (A), Firmicute (F), Alphaproteobacteria (α), Betaproteobacteria (β), Epsilonproteobacteria (ε), and Gammaproteobacteria (γ). **c** Degree of bias towards carrying VFGs more frequently than ARGs. Negative values indicate opposite bias (zero indicates no bias). Difference between empirical logits (denoted as Δempirical logit) is defined as log([*a* + 0.5]/[*b* + 0.5]) – log([*c* + 0.5]/[*d* + 0.5]), where *a* and *b* are numbers of OGVFGs and OGARGs in prophages, respectively (filled bars in b), and *c* and *d* are numbers of all OGVFGs and OGARGs in genomes, respectively (open bars in b). Numbers next to bars are P values of two-sided binomial test under null hypothesis that *a* and *b* are numbers drawn from binomial distribution with probabilities *c*/(*c* + *d*) and *d*/(*c* + *d*), respectively (corrected by Holm–Bonferroni method). Asterisks and red numbers indicate P values less than or equal to 0.05.

**Table 2.** Numbers of plasmids per genome (SD, standard deviation), and numbers of orthologous groups of antibiotic resistance genes (OGARGs) and those of virulence factor genes (OGVFGs) in genomes (total) and plasmids.

|  | Plasmid (SD) | OGARG |  | OGVFG |  |
| --- | --- | --- | --- | --- | --- |
|  |  | total | plasmid | total | plasmid |
| *A. baumannii* | 1.7 (1.6) | 455 | 195 | 48 | 0 |
| *B. anthracis* | 1.5 (0.9) | 17 | 1 | 39 | 14 |
| *B. pertussis* | 0.0 (0.0) | 0 | 0 | 95 | 0 |
| *B. melitensis* | 0.0 (0.0) | 1 | 0 | 183 | 0 |
| *B. pseudomallei* | 0.1 (0.3) | 3 | 0 | 192 | 0 |
| *C. jejuni* | 0.3 (0.6) | 65 | 21 | 480 | 9 |
| *E. faecalis* | 1.4 (1.4) | 179 | 127 | 114 | 24 |
| *E. coli* | 2.0 (1.9) | 3041 | 2397 | 18019 | 2452 |
| *H. influenzae* | 0.0 (0.1) | 7 | 1 | 161 | 0 |
| *H. pylori* | 0.2 (0.4) | 6 | 0 | 285 | 0 |
| *K. pneumoniae* | 3.4 (2.2) | 1473 | 1274 | 372 | 96 |
| *L. pneumophila* | 0.3 (0.5) | 5 | 1 | 773 | 0 |
| *L. monocytogenes* | 0.2 (0.5) | 5 | 0 | 172 | 1 |
| *M. tuberculosis* | 0.0 (0.0) | 4 | 0 | 997 | 0 |
| *N. meningitidis* | 0.0 (0.0) | 4 | 0 | 123 | 0 |
| *P. aeruginosa* | 0.2 (0.6) | 543 | 128 | 526 | 3 |
| *S. enterica* | 1.1 (1.2) | 826 | 638 | 1844 | 46 |
| *S. aureus* | 0.8 (1.0) | 423 | 182 | 217 | 9 |
| *S. agalactiae* | 0.0 (0.1) | 46 | 1 | 71 | 0 |
| *S. pyogenes* | 0.0 (0.2) | 28 | 3 | 103 | 0 |
| *V. cholerae* | 0.1 (0.4) | 101 | 36 | 251 | 0 |

### 5.4 Prophage-borne VFGs are functionally more restricted than plasmid-borne VFGs

We next asked whether chromosomal prophages and plasmids differ in the functional categories of VFGs they carry. To address this question, we mapped prophage-borne and plasmid-borne OGVFGs in *E. coli* to gene symbols (i.e., abbreviated gene names) and classified their functions using VFDB, NCBI CDD, and UniProt [38, 51, 53] (Methods under “Functional classification of prophage- and plasmid-borne VFGs”). The result shows that prophage-borne VFGs are mapped to 47 unique gene symbols, whereas plasmid-borne VFGs are mapped to 186 unique gene symbols (Table 3). This result suggests that the range of functions encoded by prophage-borne VFGs is less diverse than that encoded by plasmid-borne VFGs, as described in more detail below.

**Table 3.** Numbers of VFG symbols, orthologous groups of virulence factor genes (OGVFG), and VFGs before clustering that are detected in plasmids, chromosomal prophages, and chromosomes (which may include MGEs other than plasmids and prophages, such as transposons) in *E. coli*. Columns under ‘Chromosomes’ do not include gene symbols, OGVFGs, and VFGs that are found in neither plasmids nor chromosomal prophages. ‘Unique sum’ refers to numbers of non-redundant items (some gene symbols are counted in multiple rows owing to having multiple functions).

| Classification | Plasmids |  |  | Chromosomal prophages |  |  | Chromosomes |  |  |
| --- | --- | --- | --- | --- | --- | --- | --- | --- | --- |
|  | symbols | OGVFGs | VFGs | symbols | OGVFGs | genes | symbols | OGVFGs | VFGs |
| Adherence | 66 | 233 | 434 | 2 | 43 | 271 | 22 | 126 | 3553 |
| Biofilm | 1 | 12 | 35 | 0 | 0 | 0 | 0 | 0 | 0 |
| Dispersin | 4 | 15 | 75 | 0 | 0 | 0 | 0 | 0 | 0 |
| Enzyme | 11 | 158 | 409 | 3 | 9 | 156 | 11 | 331 | 7670 |
| Exotoxin | 15 | 236 | 1286 | 7 | 154 | 1020 | 7 | 37 | 255 |
| Immune modulation | 5 | 326 | 639 | 3 | 297 | 357 | 6 | 307 | 3075 |
| Invasion | 3 | 149 | 618 | 0 | 0 | 0 | 3 | 13 | 2881 |
| Metal uptake | 12 | 389 | 1523 | 0 | 0 | 0 | 12 | 775 | 2451 |
| Motility | 1 | 12 | 35 | 0 | 0 | 0 | 0 | 0 | 0 |
| Phage-related | 1 | 1 | 17 | 6 | 930 | 7183 | 6 | 209 | 390 |
| Regulation | 7 | 77 | 192 | 1 | 1 | 1 | 4 | 120 | 4318 |
| T2SS | 1 | 2 | 2 | 0 | 0 | 0 | 1 | 17 | 914 |
| T3SS components | 26 | 59 | 1111 | 0 | 0 | 0 | 0 | 0 | 0 |
| T3SS effectors | 33 | 659 | 1428 | 22 | 140 | 1969 | 21 | 474 | 2932 |
| T4SS | 2 | 4 | 87 | 0 | 0 | 0 | 0 | 0 | 0 |
| T5SS | 7 | 63 | 322 | 0 | 0 | 0 | 3 | 34 | 102 |
| Uncategorised | 0 | 0 | 0 | 3 | 6 | 8 | 3 | 1418 | 4557 |
| Unique sum | 186 | 2342 | 8020 | 47 | 1580 | 10965 | 97 | 3681 | 31705 |

Nearly half (viz., 22 out of 47) of the gene symbols assigned to prophage-borne VFGs (prophage-borne VFG symbols, for short) are categorised as the effectors of type III secretion system (T3SS), the proteins secreted by the T3SS apparatus (Table 3). The T3SS effectors are involved in the destruction of host cells or the modulation of host immune systems [58]. The other prophage-borne VFG symbols include those encoding exotoxins (e.g., Shiga toxins), immune modulation, and phage-related components (e.g., phage tails) (Table 3). Phage-related components are represented by the largest number of prophage-borne OGVFGs (viz., 930 out of 1580) (Table 3). Despite their abundance, the specific function of these genes is largely unknown, although some may be involved in intestinal colonisation [59, 60]. In addition, two prophage-borne VFG symbols are involved in adherence to host cells (Table 3). However, only one of them (porcine attaching-effacing associated protein, *paa*) is frequently located in prophages, whereas the other (*E. coli* common pilus chaperon, *yagV/ecpE*) is rarely located in prophages (Table S3). Taken together, the above results indicate that many prophage-borne VFGs are involved in causing direct damage to host cells or suppressing host immune systems, and a relatively small number of them are involved in host attachment.

By contrast, the gene symbols assigned to plasmid-borne VFGs (plasmid-borne VFG symbols) include a much wider range of functions, including T3SS effectors and apparatus, exotoxins, immune modulation, adherence and invasion to host cells, Type V secretion systems, enzymes (including exoenzymes), metal uptake, and gene regulation, anti-aggregation (dispersin), biofilm formation, motility, and so forth. The above results indicate that plasmid-borne VFGs encode a much wider range of functions than those encoded by prophage-borne VFGs.

Notwithstanding their comparatively small functional diversity, prophage-borne VFGs appear to play quantitatively no less important roles than those played by plasmid-borne VFGs in the virulence of *E. coli*. This is because the total number of prophage-borne OGVFGs is comparable to that of plasmid-borne OGVFGs (1580 versus 2342 OGVFGs; Table 3). Moreover, the total number of prophage-borne VFGs without clustering of orthologous genes is greater than that of plasmid-borne VFGs (10965 versus 8020 VFGs; Table 3).

Another notable pattern observed in the above analysis is that almost all gene symbols (220 out of 232) are associated with either prophage-borne or plasmid-borne VFGs, but not both (Table S3). This result means that prophages and plasmids carry distinct types of VFG even within the same functional categories. For example, within T3SS effectors, 22 and 32 gene symbols are associated with prophage-borne and plasmid-borne VFGs, respectively; however, only three gene symbols are associated with both prophage-borne and plasmid-borne VFGs (Table S3). This result suggests that prophages and plasmids have separate gene pools of virulence factors.

Taken together, the above results indicate that prophages and plasmids differ, not only in the functional diversity of VFGs they carry, but also in the specific types of VFG they carry within the same functional categories.

## 6. Discussion

The results presented above indicate that prophages tend to carry VFGs more frequently than ARGs in three of the 21 examined species. In most of the other species, prophage-borne ARGs and VFGs were hardly detected. In contrast, plasmids carry ARGs more frequently than VFGs in nine of the 21 examined species. In most of the other species, plasmid-borne ARGs and VFGs were barely detected. Taken together, these results indicate that if prophages and plasmids carry a sufficient number of VFGs or ARGs, they display opposite trends: prophages are biased towards carrying VFGs, whereas plasmids are biased toward carrying ARGs. Moreover, the comparison between prophage-borne and plasmid-borne VFGs shows that prophage-borne VFGs are functionally less diverse than plasmid-borne VFGs in *E. coli*. Taken together, the above results indicate that prophages and plasmids carry different types of bacterial accessory genes.

Based on the results presented above, we formulate the following hypothesis to test for the future (Figure 4). Temperate phages do not benefit much from carrying ARGs because if their hosts are in danger, they can abandon their hosts and seek new hosts by undergoing lytic replication [18–22]. In contrast, both phages and plasmids can benefit from VFGs because VFGs can accelerate bacterial replication by making bacteria exploit their hosts more aggressively [17]. In addition, our results suggest that plasmids benefit from a wider range of virulence functions than that from which phages benefit in *E. coli* (Table 3).

**Figure 4.**
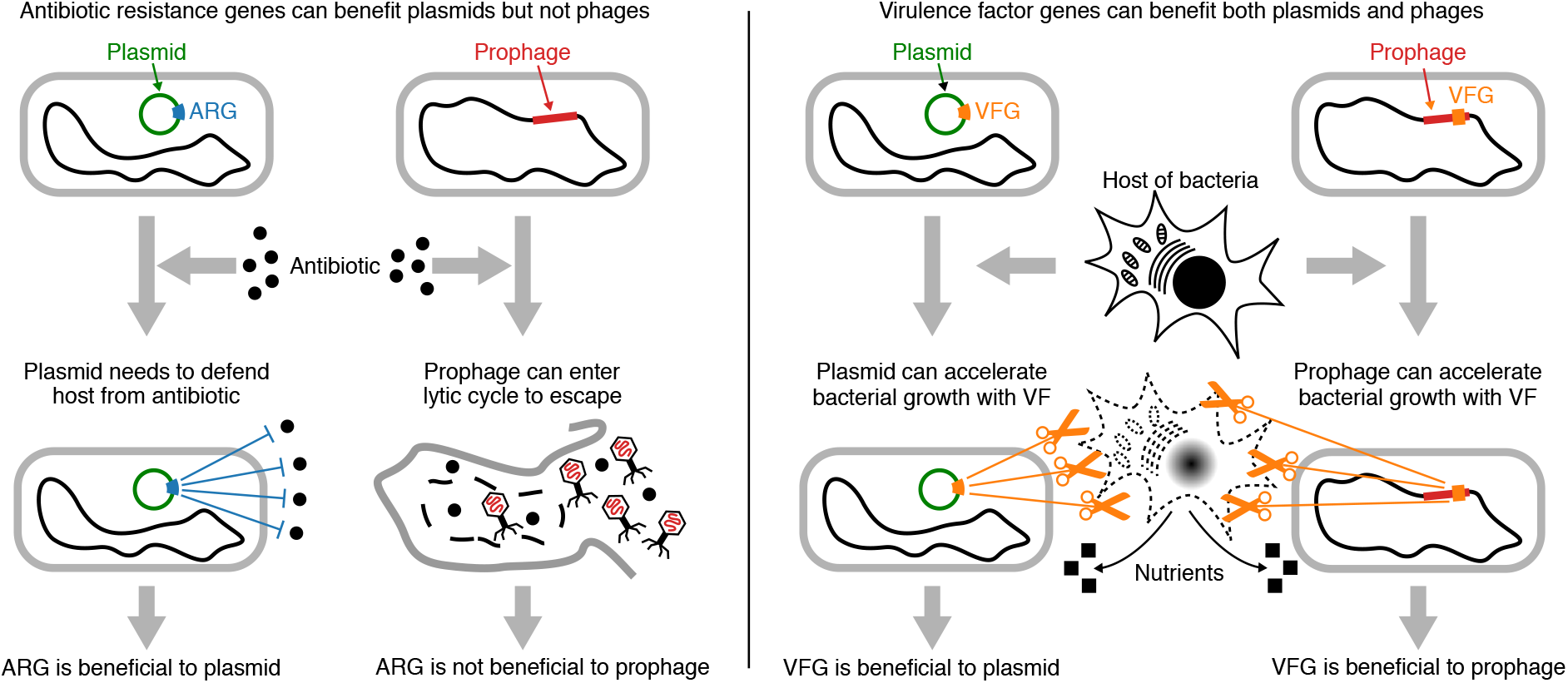
Schematic drawing of hypothesis formulated based on our results. Left: plasmids benefit from carrying ARG because they cannot abandon their hosts and thus need to minimise death of bacteria. In contrast, temperate phages do not benefit much from carrying ARGs because if their hosts are in danger, they can abandon their hosts and seek new hosts by entering lytic cycles. Right: both plasmids and phages benefit from VFGs because VFGs can enhance growth of bacteria by causing bacteria to exploit their hosts more aggressively.

The results described above, however, do not necessarily mean that plasmids rarely carry VFGs or that prophages always carry VFGs. For example, in *K. pneumoniae*, plasmids carry ARGs more frequently than VFGs; nevertheless, plasmids carry a large number of VFGs (viz., 96 out of 372 orthologous groups), whereas prophages carry none (Tables 1 and 2). A similar, yet less striking, pattern is also seen in *E. faecalis*. Given that many prophages are predicted in these species (Table 1, Figure 2), the absence of prophage-borne VFGs is unlikely to be merely due to the limited sensitivity of the prophage detection tool. By contrast, in *S. aureus* and *S. pyogenes*, prophages carry greater numbers of VFGs than do plasmids (Tables 1 and 2). This heterogeneity in the distribution of VFGs suggests that plasmids and prophages play variable roles in the pathogenicity of different bacterial species.

The absolute numbers of prophage-borne ARGs reported in this study need to be interpreted with caution because the prophage prediction tool has limited sensitivity and precision. In particular, limited sensitivity implies that prophages carrying ARGs might have been missed, so the results do not necessarily indicate that prophages are truly devoid of ARGs in many bacterial species. However, the conclusion of this study does not directly depend on the absolute numbers of prophage-borne ARGs because it is based on the comparison between the relative frequencies of prophage-borne ARGs and VFGs and those of plasmid-borne ARGs and VFGs. This comparison hinges on the assumption that the prediction of prophages carrying ARGs and VFGs are biased by common factors so that the biases are cancelled out.

Regarding the limitations of prophage prediction tools, it is pertinent to discuss a discrepancy between our result and the result of Kondo et al. (2021) with respect to *P. aeruginosa* [34]. While we found no prophage-borne ARGs in *P. aeruginosa*, Kondo et al. (2021) report that more than 10% of *P. aeruginosa* genomes possess prophage-borne ARGs [34]. The important difference between Kondo et al. (2021) and our study is that they used different tools for prophage prediction: Kondo et al. (2021) uses PHASTER [61], whereas we used VIBRANT [54]. To investigate the cause of the above discrepancy, we manually examined the 11 prophages described in Kondo et al. (2021) that carry ARGs and are predicted as ‘intact’ by PHASTER in *P. aeruginosa* [34]. We found that the examined prophages could be grouped into two categories (Table S8). In the first group (five prophages), both VIBRANT and PHASTER predicted prophages in almost the same genomic locations. However, PHASTER predicted longer genomic regions that included ARGs as prophages, whereas VIBRANT predicted shorter regions excluding ARGs. We do not know which prophage boundaries are more accurate. In the second group (six prophages), prophages were predicted only by PHASTER. These prophages, however, contained no phage-structure genes. Although they contained phage-related genes, such as integrase, transposase, and protease, these genes are not exclusively associated with phages. Moreover, PHASTER annotated tellurium resistance proteins, TerD, as virion structural proteins in three prophages in the second group, which is likely to be an error. These findings suggest that the second group of the prophages could be false positives. Prophages predicted as ‘incomplete’ or ‘questionable’ by PHASTER are less likely to be true than those predicted as ‘intact.’ Taken together, the above results suggest that the frequency of prophage-borne ARGs in *P. aeruginosa* is potentially underestimated in our study and overestimated in Kondo et al. (2021) owing to the limitations of the prophage prediction tools.

In interpreting the results obtained in this study, we assumed that ARGs and VFGs found within prophages were carried by phage genomes. However, these genes could have been inserted into pre-existing inactivated prophages (i.e., inserted after lysogeny). Although this possibility cannot be completely excluded, the following evidence suggests that not all ARGs and VFGs are inserted into pre-existing inactivated prophages. A previous study has shown that ARGs and VFGs are found in the genomes of temperate phages (which are thus not prophages) and that these genes are hardly found in the genomes of virulent phages [37]. This result would not be expected if all ARGs and VFGs were inserted into pre-existing inactivated prophages. More important, we do not have an *a priori* expectation that VFGs are more likely to be inserted into pre-existing prophages than ARGs. In the absence of such an expectation, our results are likely to be robust to post-hoc insertions of ARGs and VFGs.

That phages do not possess ARGs does not necessarily mean that phages do not mediate the horizontal transfer of ARGs because they can mediate HGT even if their genomes do not contain ARGs. Phages mediate HGT through three known mechanisms: specialised, generalised, and lateral transduction [24, 62, 63]. In specialised transduction (the focus of this study), a transferred gene constitutes a part of a phage genome [24]. By contrast, in generalised and lateral transduction, a transferred gene is originally encoded in bacterial DNA, which is encapsulated into phage particles and subsequently transferred to other cells [24, 62, 63]. Thus, phages can still mediate the horizontal transfer of ARGs even if their genomes do not contain ARGs.

In conclusion, the results presented above suggest that MGEs differ in the functional categories of accessory genes they carry depending on their strategies of infection.

## 7.3 Acknowledgements

The authors acknowledge the contribution of New Zealand eScience Infrastructure to the results of this research. The authors thank Bram van Dijk for discussion.

## 9. Supplementary information

Supplementary files can be downloaded from https://doi.org/10.1101/2022.07.21.500938

Table S1: List of all genome assemblies used in this study.

Table S2: Information about the gene-symbol assignment for OGVFGs in *Escherichia coli*, consisting of four sheets: list of VFG database entries and their gene symbols as found in the three VFG databases; manual annotation of VFG database entries that lack gene symbols, plus virB and ipgB; finalised assignment of gene symbols to VFG database entries; conflict resolution for OGVFGs that are assigned multiple gene symbols.

Table S3: List of VFG symbols, the number of OGVFG, and their functional categories.

Table S4: List of all prophages predicted by VIBRANT.

Table S5: List of all orthologous groups of antibiotic resistance genes.

Table S6: Information about all prophages manually examined.

Table S7: List of all orthologous groups of virulence factor genes.

Table S8: List of prophages that are described in Kondo et al. (2021) and manually examined in this study.

In-house scripts

