## Supplementary material for "Prophages and plasmids display opposite trends in the types of accessory genes they carry": amr-in-phage.py

```
#!/usr/bin/env python
import sys
import argparse
import re
import os

def main():
    parser = argparse.ArgumentParser(description='Check if AMR genes detected by AMRFinder reside in prophages')

    parser.add_argument('-p', '--phage_file',
                        type=argparse.FileType('r'),
                        required=True,
                        help='a TSV file that contains prophage coordinate. The format is: contig_id start end')

    parser.add_argument('-a', '--amr_file',
                        type=argparse.FileType('r'),
                        required=True,
                        help='a output file of AMRFinder')

    parser.add_argument('-o', '--out_file',
                        type=argparse.FileType('w'),
                        default=sys.stdout,
                        help='output file (default: stdout)')

    args = parser.parse_args()

    # A C-like structure
    class Bunch(object):
        def __init__(self, **kwds):
            self.__dict__.update(kwds)

    # Get phage coordinates
    phages = dict()
    for line in args.phage_file:
        words = re.split(r'\t+', line)

        contig_id = words[0]
        start = int(words[1])
        end = int(words[2])

        # If this contig_id is known
        if contig_id in phages:
            phages[contig_id].append(Bunch(start=start, end=end))

        # If this contig_id is new
        else:
            phages[contig_id] = [Bunch(start=start, end=end)]

    # Get AMR gene
    for NR, line in enumerate(args.amr_file):
        if NR > 0:
            words = re.split(r'\t+', line)
            protein_id = words[0]
            contig_id = words[1]
            start = int(words[2])
            end = int(words[3])
            comment = words[6]
            elm_type = words[8]
            elm_subtype = words[9]

            if elm_type == "AMR" and elm_subtype == "AMR":

                # The flag to tell if a gene is in a phage
                is_in_phage = False

                # If this contig has any phage 
                if contig_id in phages:
                    # Check all phages in the contig
                    for phage in phages[contig_id]:
                        # If we found a phage that includes this gene
                        if phage.start <= start and end <= phage.end:
                            is_in_phage = True

                if is_in_phage:
                    args.out_file.write('{}\t{}\tphage\t{}\t{}\t{}\n'.format(contig_id, protein_id, start, end, comment))
                else:
                    args.out_file.write('{}\t{}\tnone\t{}\t{}\t{}\n'.format(contig_id, protein_id, start, end, comment))
                    

def example(argument):
    """An example of some function."""
    return


if __name__ == '__main__':
    main()
```
