## Supplementary material for "Prophages and plasmids display opposite trends in the types of accessory genes they carry": amr-in-plasmid.py

```
#!/usr/bin/env python
import sys
import argparse
import re
import os

def main():
    parser = argparse.ArgumentParser(description='Check if AMR genes detected by AMRFinder reside in prophages')

    parser.add_argument('-p', '--plasmid_file',
                        type=argparse.FileType('r'),
                        required=True,
                        help='This should be a *_assembly_report.txt from RefSeq')
    
    parser.add_argument('-a', '--amr_file',
                        type=argparse.FileType('r'),
                        required=True,
                        help='a output file of AMRFinder')

    parser.add_argument('-o', '--out_file',
                        type=argparse.FileType('w'),
                        default=sys.stdout,
                        help='output file (default: stdout)')

    
    args = parser.parse_args()

    # A C-like structure
    class Bunch(object):
        def __init__(self, **kwds):
            self.__dict__.update(kwds)

    # Get information about contigs
    contig = dict()
    for line in args.plasmid_file:
        if line[0] != '#':
            words = re.split(r'\t+', line)
            contig_id = words[6]
            location = words[3]

            # Remember this contig
            contig[contig_id] = location

    # Get AMR gene
    for NR, line in enumerate(args.amr_file):
        if NR > 0:
            words = re.split(r'\t+', line)
            protein_id = words[0]
            contig_id = words[1]
            start = int(words[2])
            end = int(words[3])
            comment = words[6]
            elm_type = words[8]
            elm_subtype = words[9]

            if elm_type == "AMR" and elm_subtype == "AMR":
                if contig_id in contig:
                    args.out_file.write('\t'.join([contig_id, protein_id, contig[contig_id], str(start), str(end), comment]) + '\n')
                else:
                    args.out_file.write('\t'.join([contig_id, protein_id, 'ERROR', str(start), str(end), comment]) + '\n')
        
        

def example(argument):
    """An example of some function."""
    return


if __name__ == '__main__':
    main()
```
