## Supplementary material for "Prophages and plasmids display opposite trends in the types of accessory genes they carry": check-synteny-bbh.py

```
#!/usr/bin/env python
import ftplib
import urllib
import sys
import argparse
import re
import subprocess
import os
import glob
import time
import shutil
import itertools
import tempfile
import textwrap

def circle_slice(mylist, location, length):
    """
    Takes a list and return a slice as if the list is circular, where the slice consists of [location] and additional 'length' elements on both sides. In the simplest case, the slice will be [location - length: location + length + 1].
    """
    result = []
    if location-length < 0:
        result = mylist[location-length:] + mylist[0:location]
    else:
        result = mylist[location-length:location]
        
    if location + length + 1 > len(mylist):
        result = result + mylist[location:] + mylist[0:(location + length + 1) % len(mylist)]
    else:
        result = result + mylist[location:location + length + 1]
        
    return(result)

def main():
    parser = argparse.ArgumentParser(description='Ancestor tracking')

    parser.add_argument('--fasta1',
                        type=argparse.FileType('r'),
                        required=True,
                        help='This should be one of the *_translated_cds.faa files')

    parser.add_argument('--fasta2',
                        type=argparse.FileType('r'),
                        required=True,
                        help='This argument should be one of the 2 *_translated_cds.faa file')
    
    parser.add_argument('-o', '--output_file',
                        type=argparse.FileType('w'),
                        default=sys.stdout,
                        help='output file (default: stdout)')
    
    parser.add_argument('-n', '--neighbor-distance',
                        type=int,
                        default=3,
                        help="synteny is checked for a region consisting of '1 + 2*neighbor-distance' genes, namely, a focal gene and 'neighbor-distance' genes upstream or downstream to the focal gene (default: %(default)s)")
    
    parser.add_argument('file',
                        nargs = '?',
                        type=argparse.FileType('r'),
                        default=sys.stdin,
                        help='This is a file containing the output of proteinortho Step 2 (blast-graph). The file should contain all the network information about the two genomes provided by *_translated_cds.faa files (see [files] argument). It may contain network information of other genome pairs. (default: stdin)')

    args = parser.parse_args()

    # A C-like structure
    class Bunch(object):
        def __init__(self, **kwds):
            self.__dict__.update(kwds)

    # Exist if two files are identical
    if args.fasta1.name == args.fasta2.name:
        sys.stderr.write('ERROR: Two input files are identical\n')
        exit(1)
            
    # The next variable ('genomes') reads args.fasta1. 'genomes' is a
    # list of two dictionaries, each reads either args.fasta1 or
    # args.fasta2. These dictionaries map a contig name to a list of
    # genes, in which the genes are ordered in the way as in the genome.
    genomes = [dict(), dict()]

    # Next variable ('gene_to_location') contains the information about
    # the location of genes. 'gene_to_location' is a dictionary that
    # maps a gene name to a tuple of three elements:
    # [0] an integer indicating the genome
    # [1] the name of the contig
    # [2] the index of this gene in the list that is contained in 'genomes'.
    gene_to_location = dict()

    # Regular expression to get the contig name
    re_contig = re.compile('^>lcl[|](.._[^_]+)_')

    # Regular expression to get the gene name
    re_gene = re.compile('^>(lcl[^ ]+) ')

    # Read genomes and synteny information
    for genome_index, fin in enumerate([args.fasta1, args.fasta2]):
        for line in fin:
            # Read protein name
            if line[0] == '>':
                contig = re_contig.match(line).group(1)
                gene = re_gene.match(line).group(1)

                # If this contig hasn't been created
                if not contig in genomes[genome_index]:
                    genomes[genome_index][contig] = [gene]
                    gene_to_location[gene] = (genome_index, contig, len(genomes[genome_index][contig])-1)
                else:
                    genomes[genome_index][contig].append(gene)
                    gene_to_location[gene] = (genome_index, contig, len(genomes[genome_index][contig])-1)

    
                    
    # This will map a gene to the genes to which the first gene is connected
    network_edges = dict()

    # This will map a pair of genes to the corresponding network
    # information in args.file. This will be used when we
    # generate output.
    gene_pair_to_network_info = dict()

    # This contains header of args.file. This will be used when
    # we generate output.
    header = []

    # Flag indicating that this part of args.file is about args.fasta1.name and args.fasta2.name
    this_is_correct_genome_pair = False

    # This is a set of the fasta file names
    fasta_file_names = set([os.path.basename(args.fasta1.name), os.path.basename(args.fasta2.name)])
    
    # Read network information
    for line in args.file:

        # If this line is a comment
        if line[0] == '#':
            # Check if this comment is about args.fasta1.name and args.fasta2.name
            if fasta_file_names == set(line[1:].strip().split()):
                # We will start to read the file
                this_is_correct_genome_pair = True

                # Remember comment
                header.append(line)
                
            # If we have finished reading all network edges between
            # args.fasta1.name and args.fasta2.name. 
            elif this_is_correct_genome_pair and network_edges:
                # We have finished reading the relevant part.
                break
            # If we have started to read network edges, but have not
            # read any of them
            elif  this_is_correct_genome_pair:
                # Remember comment
                header.append(line)

            # If we have not started to read network edges
            else:
                # Remember comment if need be
                if len(header) < 2:
                    header.append(line)

            # This is a comment line, so skip it.
            continue

        # If this line is not a comment and if we started to read network edges
        elif this_is_correct_genome_pair:
            words = line.split()

            # Add this edge to network_edges
            if not words[0] in network_edges:
                network_edges[words[0]] = [words[1]]
            else:
                network_edges[words[0]].append(words[1])

            # Add this edge from the opposite side too
            if not words[1] in network_edges:
                network_edges[words[1]] = [words[0]]
            else:
                network_edges[words[1]].append(words[0])

            # Remember this edge
            gene_pair_to_network_info[frozenset(words[0:2])] = line

    # Output header
    args.output_file.write(''.join(header))

    # Go through every gene in one genome and output the network information
    for gene_x in itertools.chain(*genomes[0].values()):
        # Check if gene_x has any homolgs in the other genome.
        if not gene_x in network_edges:
            # If it has no homolog, go to the next gene.
            continue

        # Get the homologs of gene_x
        list_of_gene_y = network_edges[gene_x]
            
        # Get the location of gene_x
        genome_index, contig, gene_index = gene_to_location[gene_x]

        # Get a genomic region around gene_x
        region_around_gene_x = circle_slice(genomes[genome_index][contig], gene_index, args.neighbor_distance)

 
        # The next variable will get the homologs of the genes in
        # region_around_gene_x.
        homologs_of_region_around_gene_x = set()
        for gene_near_gene_x in region_around_gene_x:
            if gene_near_gene_x in network_edges:
                homologs_of_region_around_gene_x.update(network_edges[gene_near_gene_x])

        # Go through all the homologs of gene_x
        for gene_y in list_of_gene_y:
            # Error check
            if not gene_y in gene_to_location:
                sys.stderr.write('gene_x is '+gene_x+'\n')

            # Get the location of gene_y
            genome_index, contig, gene_index = gene_to_location[gene_y]

            # Get a genomic region around gene_y
            region_around_gene_y = circle_slice(genomes[genome_index][contig], gene_index, args.neighbor_distance)

            # Count how many homologs of the genes in
            # region_around_gene_x are found in region_around_gene_y
            n_of_homologs_of_region_around_gene_x_in_region_around_gene_y = len(homologs_of_region_around_gene_x.intersection(region_around_gene_y))

            # The next variable will get the homologs of the genes in
            # region_around_gene_y.
            homologs_of_region_around_gene_y = set()
            for gene_near_gene_y in region_around_gene_y:
                if gene_near_gene_y in network_edges:
                    homologs_of_region_around_gene_y.update(network_edges[gene_near_gene_y])

            # Count how many homologs of the genes in
            # region_around_gene_y are found in region_around_gene_x
            n_of_homologs_of_region_around_gene_y_in_region_around_gene_x = len(homologs_of_region_around_gene_y.intersection(region_around_gene_x))

            # If this edge is supported by synteny, output it.
            if args.neighbor_distance < min(n_of_homologs_of_region_around_gene_x_in_region_around_gene_y,n_of_homologs_of_region_around_gene_y_in_region_around_gene_x):
                if frozenset([gene_x,gene_y]) in gene_pair_to_network_info:
                    args.output_file.write(gene_pair_to_network_info[frozenset([gene_x,gene_y])])
                else:
                    sys.stderr.write('ERROR: {} does not contain gene pair, {} and {}\n'.format(args.file.name, gene_x, gene_y))
                    exit(1)
                
def example(argument):
    """An example of some function."""
    return


if __name__ == '__main__':
    main()
```
