## Supplementary material for "Prophages and plasmids display opposite trends in the types of accessory genes they carry": cluster-genes.py

```
#!/usr/bin/env python
import sys
import argparse
import re
import os

def main():
    parser = argparse.ArgumentParser(description='Cluster virulence or AMR genes according to the clusters generated by proteinortho. It takes a file created by merge-phage-plasmid.py and creates files, each of which contains one cluster that contains ARG or VFG.')

    parser.add_argument('-p', '--proteinortho_file',
                        type=argparse.FileType('r'),
                        required=True,
                        help='The tsv file (myproject.proteinortho.tsv) generated by proteinortho.')
    
    # parser.add_argument('-a', '--amr_file',
    #                     type=argparse.FileType('r'),
    #                     required=True,
    #                     help='a output file of AMRFinder')

    parser.add_argument('-o', '--out_prefix',
                        default='cluster_',
                        help='prefix of output file (default: %(default)s)')

    parser.add_argument('-x', '--gene_postfix',
                        required=True,
                        help='the postfix of file names containing virulence or AMR genes (see below). The postfix will be removed from file names to create assembly names.')
    
    parser.add_argument('-y', '--genome_postfix',
                        required=True,
                        default='_translated_cds.faa',
                        help='the postfix of NCBI file names containing genome data (see below). The postfix will be removed from file names to create assembly names.  (default: %(default)s)')

    parser.add_argument('infile',
                        nargs = '?',
                        type=argparse.FileType('r'),
                        default=sys.stdin,
                        help='path to a file containing virulence or AMR genes that are output of merge-phage-plasmid.py (default: stdin).')
    
    args = parser.parse_args()
 
    # A C-like structure
    class Bunch(object):
        def __init__(self, **kwds):
            self.__dict__.update(kwds)

    all_genes = dict()
            
    # Get information about virulence or AMR genes in args.files
    for line in args.infile:
        words = re.split(r'\t+', line.strip())

        assembly = words[0]
        contig = words[1]
        protein = words[2]
        plasmid = words[3]
        phage = words[4]
        start = words[5]
        end = words[6]
        comment = words[-1]

        # remember this gene
        if (assembly, protein) in all_genes:
            all_genes[(assembly, protein)].append(line.strip())
        else:
            all_genes[(assembly, protein)] = [line.strip()]

    # This will get the names of all assemblies.
    assemblies = []

    # Go through all clusters. Each line describes one cluster, except
    # the 0th line, which contains assembly names.
    for NR, line in enumerate(args.proteinortho_file):
        # Get the assembly name
        if NR == 0:
            words = re.split(r'\t+', line.strip())
            for word in words[3:]:
                n = word.find(args.genome_postfix)
                if n == -1:
                    sys.stderr.write('ERROR: {} has no postfix\n'.format(fname))
                    exit(1)
                else:
                    assemblies.append(word[:n])

            # Go to the next line
            continue


        # If NR>1, get a cluster
        words = re.split(r'\t+', line.strip())
        
        # This will get information about one cluster. 
        cluster = []

        # NF (column index) tells which assembly this column is. word
        # contains a comma-separated list of proteins
        for NF, word in enumerate(words[3:]):
            if word != '*':
                proteins = re.split(r',', word)
                for protein in proteins:
                    # The taple used in the next line is also used as a
                    # key of 'all_genes', which is a dictionary
                    # containing all virulence or AMR genes.
                    cluster.append((assemblies[NF], protein))

        # Output cluster if it contains virulence genes
        output = []
        for key in cluster:
            
            # If this 'key' is a virulence or AMR gene
            if key in all_genes:
                # Add it to the output line
                output.extend(all_genes[key])

        if output:
            with open('{}{:06d}.tsv'.format(args.out_prefix, NR), 'w') as fout:
                for line in output:
                    fout.write(line+'\n')
            
    # contig = dict()
    #     if line[0] != '#':
    #         words = re.split(r'\t+', line)
    #         contig_id = words[6]
    #         location = words[3]

    #         # Remember this contig
    #         contig[contig_id] = location


    #         if elm_type == "AMR" and elm_subtype == "AMR":
    #             if contig_id in contig:
    #                 args.out_file.write('{}\t{}\t{}\t{}\n'.format(contig_id, protein_id, contig[contig_id], comment))
    #             else:
    #                 args.out_file.write('{}\t{}\tERROR\t{}\n'.format(contig_id, protein_id, comment))
        
        

def example(argument):
    """An example of some function."""
    return


if __name__ == '__main__':
    main()
```
