## Supplementary material for "Prophages and plasmids display opposite trends in the types of accessory genes they carry": merge-phage-plasmid.py

```
#!/usr/bin/env python
import sys
import argparse
import re
import os

def main():
    parser = argparse.ArgumentParser(description='This takes a file created by virulence-in-plasmid.py or amr-in-plasmid.py and creates a file that should be given to cluster-genes.py.')

    parser.add_argument('-v', '--phage_file',
                        required=True,
                        help='an outout file of virulence-in-phage.py or amr-in-phage.py')
    
    parser.add_argument('-o', '--out_file',
                        type=argparse.FileType('w'),
                        default=sys.stdout,
                        help='output file (default: stdout)')

    parser.add_argument('-x', '--postfix',
                        required=True,
                        help='the postfix of plasmid_file. The postfix will be removed from the file name to create the assembly name.')

    parser.add_argument('plasmid_file',
                        nargs = 1,
                        #nargs = 2 # if there should be 2 files
                        type=str,
                        help='an output file of virulence-in-plasmid.py or amr-in-plasmid.py')

    args = parser.parse_args()
 
    # A C-like structure
    class Bunch(object):
        def __init__(self, **kwds):
            self.__dict__.update(kwds)

    all_genes = dict()
            
    # Get the assembly name from the file name
    assembly = os.path.basename(args.plasmid_file[0])
    n = assembly.find(args.postfix)

    if n == -1:
        sys.stderr.write('ERROR: {} has no postfix\n'.format(args.plasmid_file))
        exit(1)
    else:
        assembly = assembly[:n]

    # Get information about the prophage-context of genes
    with open(args.phage_file) as fin:
        for line in fin:
            words = re.split(r'\t+', line.strip())

            contig = words[0]
            protein = words[1]
            phage = words[2]
            start = words[3]
            end = words[4]
            comment = words[-1]

            # remember this gene
            all_genes[(assembly, protein, start, end)] = [assembly, contig, protein, phage, start, end, comment]

    # Get information about the plasmid-context of genes
    with open(args.plasmid_file[0]) as fin:
        for line in fin:
            words = re.split(r'\t+', line.strip())

            contig = words[0]
            protein = words[1]
            plasmid = words[2]
            start = words[3]
            end = words[4]
            comment = words[-1]

            # Check if this gene has been recorded
            if (assembly, protein, start, end) in all_genes:
                all_genes[(assembly, protein, start, end)].insert(3, plasmid)
            else:
                sys.stderr.write('ERROR: {} in {} is not found in {}\n'.format(protein, args.phage_file, args.plasmid_file[0]))

    # Output poteins
    for key, value in all_genes.items():
        args.out_file.write('\t'.join(value) + '\n')
        
        

def example(argument):
    """An example of some function."""
    return


if __name__ == '__main__':
    main()
```
