## Supplementary material for "Prophages and plasmids display opposite trends in the types of accessory genes they carry": README

check-synteny-bbh.py: Parse the output of the step 2 of ProteinOrtho and
produce the input for the step 3 of ProteinOrtho by pruning reciprocal
nearly-best hits produced by ProteinOrtho based on synteny information.

arm-in-plasmid.py: Parse the output of NCBI AMRFinderPlus and an RefSeq
assembly report file and produce a list of ARGs in plasmids.

amr-in-phage.py: Parse the output of NCBI AMRFinderPlus and a file
containing prophage coordinates and produce a list of ARGs in prophages.

select-best-hit-cds.py: Parse the output of BLASTP with multiple VFG
queries to find the best hit VFG.

virulence-in-phage.py: Parse the output of select-best-hit-cds.py to
select genes encoded in prophages that meet the three criteria to be
considered as encoding VF: percent identity, coverate, and species name.

virulence-in-plasmid.py: Parse the output of select-best-hit-cds.py to
select genes encoded in plasmids that meet the three criteria to be
considered as encoding VF: percent identity, coverate, and species name.

merge-phage-plasmid.py: Parse the output of arm-in-plasmid.py and
arm-in-phage.py (or virulence-in-plasmid.py and virulence-in-phage.py)
to merge information about whether ARGs (or VFGs) are encoded in
plasmids or prophages.

cluster-genes.py: Parse the output of the step 3 of ProteinOrtho and
merge-phage-plasmid.py and create files, each of which contains one
orthologous cluster of VFGs (or ARGs).
