## Supplementary material for "Prophages and plasmids display opposite trends in the types of accessory genes they carry": select-best-hit-cds.py

```
#!/usr/bin/env python
import sys
import argparse
import re
import os

def main():
    parser = argparse.ArgumentParser(description='Multiple virulence factor genes can hit the same gene in a bacterial genome. This program selects the virulence factor gene that has the best e-value to a gene in a bacterial genome. This script assumes that blast databases are created from *_translated_cds.faa. ')

    parser.add_argument('-b', '--blast_file',
                        type=argparse.FileType('r'),
                        required=True,
                        help='an output file of blastp')

    parser.add_argument('-f', '--faa_file',
                        type=argparse.FileType('r'),
                        required=True,
                        help='a *_translated_cds.faa file of RefSeq')

    parser.add_argument('-o', '--out_file',
                        type=argparse.FileType('w'),
                        default=sys.stdout,
                        help='output file (default: stdout)')

    
    args = parser.parse_args()

    # A C-like structure
    class Bunch(object):
        def __init__(self, **kwds):
            self.__dict__.update(kwds)

    # This dictionary maps the gene_id of a blast hit to the various
    # information about the blast hit such as e-value, % identity,
    # contig name, coordinate in the contic, etc.
    hit = dict()

    # Read blast output file and get hits.
    blast_file = iter(args.blast_file)
    for line in blast_file:
        # If this is the begining of new result entry
        if line[0:8] == '# BLASTP':
            # Get the name of the query gene
            line = next(blast_file)
            query = ' '.join(line.strip().split()[2:])
        # Skip comment lines
        elif line[0] == '#':
            pass
        else:
            words = re.split(r'\t+', line)
            gene_id = words[0]
            evalue = float(words[1])
            bitscore = float(words[2])
            length = int(words[3])
            slen = int(words[4])
            sstart = int(words[5])
            send = int(words[6])
            qlen = int(words[7])
            qstart = int(words[8])
            qend = int(words[9])
            pindent = float(words[10])
            mincov = 100*min((send-sstart)/slen, (qend-qstart)/qlen)

            # If this gene has been included in 'hit'
            if gene_id in hit:
                # If this is a better hit, replace the worse hit with it.
                if hit[gene_id].bitscore < bitscore:
                    hit[gene_id] = Bunch(evalue=evalue, bitscore=bitscore, pindent=pindent, mincov=mincov, query=query, contig_id=None, coord=None)
            # If this gene has not been included in 'hit'
            else:
                # Add this gene to 'hit'
                hit[gene_id] = Bunch(evalue=evalue, bitscore=bitscore, pindent=pindent, mincov=mincov, query=query, contig_id=None, coord=None)

    # Regular expression to get the gene name
    re_gene = re.compile('^>(lcl[^ ]+) ')

    # Regular expression to get the contig name of a gene in a FAA file
    re_contig = re.compile('^>lcl[|](.._[^_]+)_')

    # Regular expression to get the location field of a gene in a FAA file
    re_location = re.compile('\[location=([^]]+)\]')

    # Regular expression to get the coordinates within a location field
    re_number = re.compile('[0-9]+')
    
    # Read a FAA file and get the contig_id and coordinate of each hit
    find_name = re.compile('Name=([^;]+);')
    for line in args.faa_file:
        if line[0] == '>':
            # Get the name of the gene (CDS)
            gene_id = re_gene.search(line).group(1)
            
            # If this gene is in the list of hits
            if gene_id in hit:
                # Get the name of the contig containing this gene, and
                # the coordinates of this gene in the contic,
                contig_id = re_contig.search(line).group(1)
                location = re_location.search(line).group(1)
                coordinate = re_number.findall(location)
                
                # Add the coordinate and contig_id
                if hit[gene_id].contig_id == None:
                    hit[gene_id].contig_id = [contig_id]
                    hit[gene_id].coord = [(coordinate[0], coordinate[-1])]
                # It's impossible that the same gene appears multiple times
                else:
                    sys.stderr.write('ERROR: the same gene appears multiple times in the FAA file\n')
                
    args.out_file.write('#subject id, contig, start, end, evalue, % identity, % coverage, virulence\n')
    for key, value in hit.items():
        if value.contig_id == None:
            sys.stderr.write('ERROR: {} is missing in GFF\n'.format(key))
            continue

        for i in range(len(value.contig_id)):
            args.out_file.write('\t'.join([key, value.contig_id[i], value.coord[i][0], value.coord[i][1], '{:.02e}'.format(value.evalue), '{:.03f}'.format(value.pindent), '{:.03f}'.format(value.mincov), value.query]) + '\n')
        

def example(argument):
    """An example of some function."""
    return


if __name__ == '__main__':
    main()
```
