## Supplementary material for "Prophages and plasmids display opposite trends in the types of accessory genes they carry": virulence-in-phage.py

```
#!/usr/bin/env python
import sys
import argparse
import re
import os

def main():
    parser = argparse.ArgumentParser(description='Check if virulence genes reside in prophages')

    parser.add_argument('-p', '--phage_file',
                        type=argparse.FileType('r'),
                        required=True,
                        help='a TSV file that contains prophage coordinate. The format is: contig_id start end')
    
    parser.add_argument('-v', '--virulence_file',
                        type=argparse.FileType('r'),
                        required=True,
                        help='a output file of virulence-blast.py')

    parser.add_argument('-o', '--out_file',
                        type=argparse.FileType('w'),
                        default=sys.stdout,
                        help='output file (default: stdout)')

    parser.add_argument('-e', '--evalue',
                        type=float,
                        default=1e-9,
                        help='minimum evalue (default: %(default)s)')

    parser.add_argument('-c', '--coverage',
                        type=float,
                        default=80.0,
                        help='minimum percent alignment coverage (default: %(default)s)')

    parser.add_argument('-i', '--identity',
                        type=float,
                        default=80.0,
                        help='minimum percent sequence identity (default: %(default)s)')

    parser.add_argument('-s', '--species',
                        default='all',
                        help='Only consider virulence genes of the specified species. For example, --species \'Salmonella enterica\' will take only genes from Salmonella enterica (--species \'Escherichia coli\' is an exception, in that it also includes Shigella sp.). Setting --species \'all\' will include all species. (default: %(default)s)')

    
    args = parser.parse_args()

    # A C-like structure
    class Bunch(object):
        def __init__(self, **kwds):
            self.__dict__.update(kwds)

    # Get the name of species to consider
    species_included = set()
    if args.species != 'all':
        if args.species == 'Escherichia coli':
            species_included.add('Escherichia coli')
            species_included.add('Shigella boydii')
            species_included.add('Shigella sonnei')
            species_included.add('Shigella dysenteriae')
            species_included.add('Shigella flexneri')
        else:
            species_included.add(args.species)
            
    # Get phage coordinates
    phages = dict()
    for line in args.phage_file:
        words = re.split(r'\t+', line)
        contig_id = words[0]
        start = int(words[1])
        end = int(words[2])
        
        # If this contig_id is known
        if contig_id in phages:
            phages[contig_id].append(Bunch(start=start, end=end))

        # If this contig_id is new
        else:
            phages[contig_id] = [Bunch(start=start, end=end)]

    # Get virulence gene
    get_species = re.compile('\[([^]]+)\]$')
    for line in args.virulence_file:
        if line[0] == '#':
            continue
        
        words = re.split(r'\t+', line)
        protein_id = words[0]
        contig_id = words[1]
        start = int(words[2])
        end = int(words[3])
        evalue = float(words[4])
        identity = float(words[5])
        coverage = float(words[6])
        comment = words[7].strip()

        species = ' '.join(get_species.search(comment).group(1).split()[0:2])

        if (evalue <= args.evalue and
            identity >= args.identity and
            coverage >= args.coverage and
            (species in species_included or args.species=='all')):


            # The flag to tell if a gene is in a phage
            is_in_phage = False

            # If this contig has any phage 
            if contig_id in phages:
                # Check all phages in the contig
                for phage in phages[contig_id]:
                    # If we found a phage that includes this gene
                    if phage.start <= start and end <= phage.end:
                        is_in_phage = True
            
            if is_in_phage:
                args.out_file.write('\t'.join([contig_id, protein_id, 'phage', str(start), str(end), comment]) + '\n')
            else:
                args.out_file.write('\t'.join([contig_id, protein_id, 'none', str(start), str(end), comment]) + '\n')
        

def example(argument):
    """An example of some function."""
    return


if __name__ == '__main__':
    main()
```
